# Dorsal Hippocampus Not Always Necessary in a Radial Arm Maze Delayed Win-shift Task

**DOI:** 10.1101/433565

**Authors:** Dylan Layfield, Nathan Sidell, Afnan Abdullahi, Ehren L. Newman

## Abstract

Spatial working memory is important for foraging and navigating the environment. However, its neural underpinnings remain poorly understood. The hippocampus, known for its spatial coding and involvement in spatial memory, is widely understood to be necessary for spatial working memory when retention intervals increase beyond seconds into minutes. Here, we describe new evidence that the dorsal hippocampus is not always necessary for spatial working memory for retention intervals of 8 minutes. Rats were trained to perform a delayed spatial win shift radial arm maze task (DSWS) with an 8-minute delay between study and test phases. We then tested whether bilateral inactivation of the dorsal hippocampus between the study and test phases impaired behavioral performance at test. Inactivation was achieved through a bilateral infusion of lidocaine. Performance following lidocaine was compared to control trials, in which, sterile phosphate buffered saline (PBS) was infused. Test performance did not differ between the lidocaine and PBS conditions, remaining high in each. To explore the possibility that this insensitivity to inactivation was a result of overtraining, a second cohort of animals received substantially less training prior to the infusions. In this second cohort, lidocaine infusions did significantly impair task performance. These data indicate that successful performance of a spatial win-shift task on the 8-arm maze need not always be hippocampally dependent.

## 1. Introduction

The ability to forage is key to survival for many animals. A critical component of foraging behavior is spatial working memory, wherein information regarding the spatial positions that food has been found or that remain to be searched is maintained (Olton & Samuelson, 1976). Identifying the brain structures that support spatial working memory remains an outstanding goal of behavioral neuroscience. Pharmacological inactivation and lesion studies indicate that the hippocampus is necessary when the delay between encoding of spatial information and use of that information increases from seconds to minutes (Lee & Kesner., 2003a; Lee & Kesner., 2003b; Churchwell & Kesner., 2011).

Spatial working memory in rodents is often studied using the radial arm maze (Olton & Samuelson., 1976). A widely used radial arm maze task is the delayed spatial win-shift task (DSWS; Packard et al., 1990; Seamans and Phillips., 1994; Seamans et al., 1995). In this task, rats first complete a study phase with a subset of arms available and baited with food. Later, rats complete a test phase where all arms are open, and the rat is rewarded for visiting previously un-entered arms. Though prior work has shown that inactivation of the ventral hippocampus/subiculum reduced performance after a 30-minute delay (Floresco et al., 1997), the necessity of the dorsal hippocampus for this task remains unknown.

Given the existing literature showing the hippocampus is involved in spatial cognition (O’Keefe & Nadel., 1978; Buzsaki & Moser., 2013; Hartley et al., 2014) and that lesions / pharmacological inactivations of dorsal hippocampus impair performance in other spatial working memory tasks (McDaniel et al., 1994; Lee & Kesner., 2003a; Lee & Kesner, 2003b; Potvin et al., 2006; Yoon et al., 2008), we expected that the dorsal hippocampus would be necessary for accurate test performance the DSWS task on the radial arm maze with 8 min retention intervals. However, here, we describe data showing that bilateral infusions of lidocaine administered at the outset of an 8-minute retention interval between study and test did not impair performance relative to phosphate buffered saline infusions. To test if this insensitivity was possibly the result of overtraining, a second cohort of animals received substantially less training prior to the infusions. In this second cohort, lidocaine infusions significantly impaired task performance. These results challenge the traditional view that the dorsal hippocampus is always needed for spatial memory guided behavior at intermediate delays.

## 2. Methods

The data presented here was collected incidentally in the running of a larger study. In the broader study, the trials of interest were those with delays of 60 min or longer. Trials with 8 min retention intervals had been included to control for expectancy effects central to that design. We describe here the full set of methods but will focus our analyses on short retention interval trials, during which, the inactivation could be expected to be in effect.

All animal procedures and surgery were conducted in strict accordance with National Institutes of Health the Indiana University Institutional Animal Care and Use Committee guidelines.

### 2.1 Subjects

19 Male Adult Long Evans rats were used: 10 in cohort one and nine in cohort two. One animal was dropped from cohort one due to inaccurate cannula placement. Rats were individually housed and maintained on a 12 H light/dark cycle in a temperature and humidity-controlled room with ad libitum access to water, and food restricted to maintain ~90% (85-95%) of free feeding body weight. Rats were acclimated to the animal facility for 5 days before being handled daily.

### 2.2 Behavioral training

Training took place on a custom automated radial maze (Maze Engineers, Cambridge, MA) with a 33.2 cm wide hub and pneumatic drop doors at the entrances to the 8 arms, each measuring 48.26 cm long, 10.79 cm wide with 20.95 cm tall walls. At the end of each arm were food wells in which 45 mg sucrose pellets (Bio-Serv, Flemington, NJ) were delivered. The maze was open on top, with clear acrylic walls, allowing for viewing of a variety of distal cues surround the maze. The maze was cleaned with chlorohexadine immediately after each trial. Figure 1A summarizes the training and testing phases rats in cohort one and two received.

**Fig 1.**
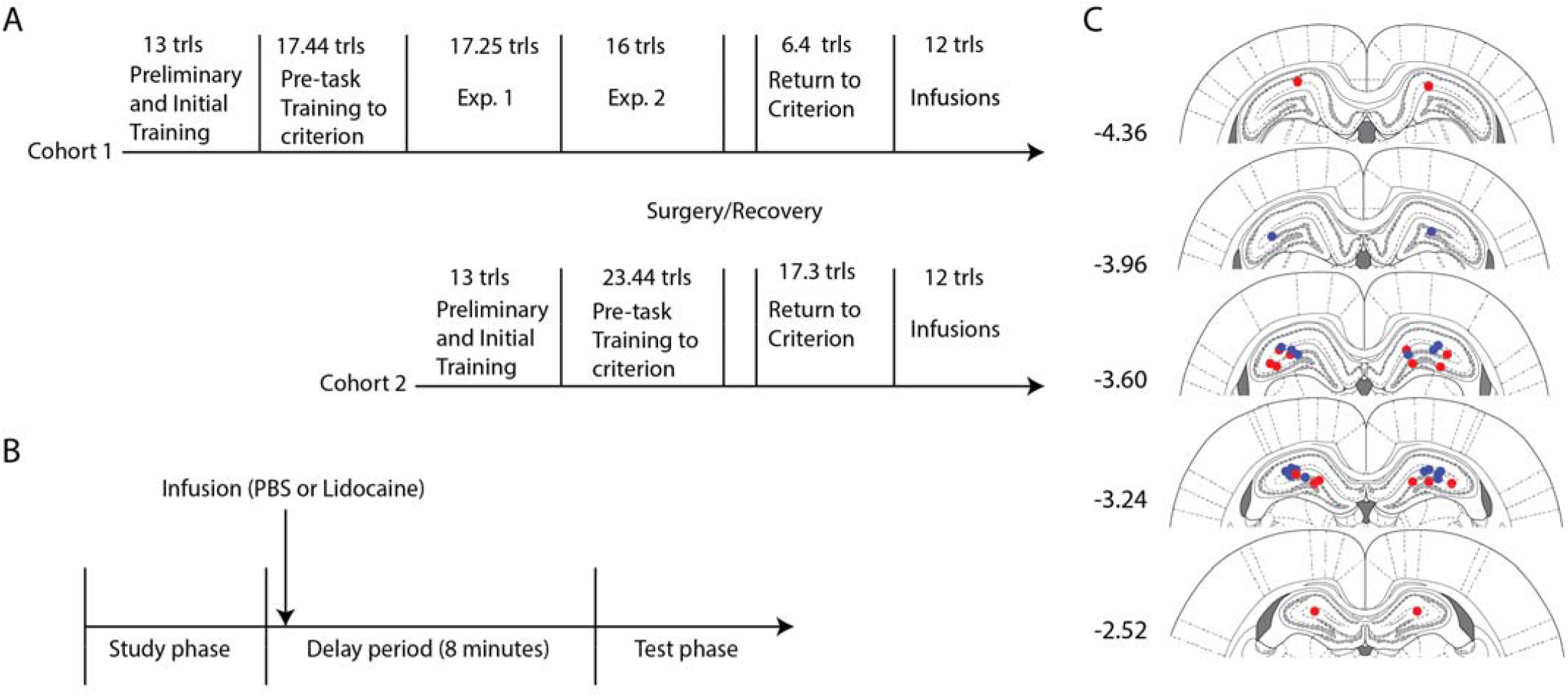
Experiment overview. (A) Timeline of task training. Rats in cohort one and two received the same training up to reaching behavioral criterion. Only cohort one participated in two pilot studies prior to surgery. Both cohorts received the same training following surgery. The mean number of trials required at each step is shown. (B) Timeline of task trials. Rats were placed in the hub for 1 minute before 4 doors open in the study phase. Immediately following study phase rats were given infusions and placed back in their home cage (in the maze room) for remainder of the delay. After the 8-minute delay, rats were placed back in the hub for 1 minute before all 8 doors open for test phase. (C) Coronal plates showing cannula termination spots. Each circle represents a cannula termination spot for a rat. (Red circles cohort 1, Blue circles cohort 2). Numbers next to each plate indicate distance from bregma in mm. trls = trials.

#### 2.2.1 Habituation & Preliminary Training

For 3 days prior to training, rats were handled for 10 minutes and given 20-30 sucrose pellets to habituate them to experimenter handing and rewards. Preliminary training consisted of one 10 min trial daily for 4 days. During each, 4-6 pellets were placed along each of the 8 arms and 2 pellets in the food wells of each arm. The trial ended after the rat consumed all pellets or 10 minutes had elapsed.

#### 2.2.2 Initial Training

Initial Training consisted of 10 sessions, 1 trial/session. During each, rats were trained that arms would be baited once per day with 2 pellets. Training trials began by placing the animal in the central hub with testing room lights off and hub doors closed. After 1-min the doors opened automatically, and lights turned on allowing the rat to freely forage. Lights were turned off during all hub placements in this and all future phases of training to prevent increased exposure to extra-maze cues. The rats could explore until all pellets were collected or 15 minutes had elapsed.

#### 2.2.3 Pre-task Training

Pre-task training consisted of two phases, a study phase and a test phase. In the study phase, the rat was placed in the central hub, after 1 minute, a random set of four doors opened, and the rat was allowed to collect pellets from each. The rat was then removed and placed in their home cage for an 8-min retention phase. During which, the maze was cleaned. The test phase began by placing the rat in the hub and, after 1 min, all eight doors opened. The four arms not opened at study were baited with pellets. The test phase ended after all the pellets had been consumed or when 15 minutes elapsed. The trial timeline is illustrated in Figure 1B. Criterion performance, required before moving on in the study, was no more than three errors over four days, with any days having errors requiring the optimal number of arm entries (for example 1 error requires 5 arm entries 2 errors 6 arm entries etc.).

#### 2.2.4 Extended task experience prior to surgery

Rats in cohort one participated in two pilot experiments between reaching criterion and receiving surgery (Figure 1). These pilot experiments consisted of test runs through the task described below. These variants used retention intervals of 8, 60, and 150 minutes and varied the holding location of rats during the retention interval. Rats in cohort two did not participate in pilot studies, reducing the total experience with the task prior to surgery (see Results for comparison of training time between cohorts).

#### 2.2.5 Task

The larger task design, from which the current trials of interest were drawn, replicated the ‘retrieval practice’ design of Crystal et al. (2013). In short, rats performed a DSWS task on an 8-arm maze but on some trials, rats were placed back on the track at 8 min but removed again prior to the doors opening. The full test phase was then run after the remainder of a 60 min delay. Our study used a fully counterbalanced 3 × 2 design with 3 behavioral conditions {8 min retention, 60 min retention with retrieval practice, 60 min retention without retrieval practice} and 2 infusion types {lidocaine, phosphate buffered saline}.

### 2.3. Surgery

Rats were anesthetized with isoflurane (1 – 4% in oxygen) and placed in a stereotaxic frame (Kopf Instruments). A scalp incision was made, 2-4 screws were inserted into the skull and two craniotomies were drilled to target dorsal hippocampus with the coordinates AP: −3.8 mm, ML ±2.5 mm. Two 26-gauge guide cannulas (Plastics one) were lowered into the brain, DV 1.8 mm from brain surface, and secured to the anchor screws with dental acrylic. Dummy cannulas were inserted into the guide cannulas. Rats recovered for 5+ days post-surgery.

### 2.4. Intracranial microinfusions

For infusions, the dummy cannulas were removed and injector cannulas with 1 mm projection prefilled with solution was inserted. A microinfusion pump infused a total volume of 0.5ul at 0.5ul per minute per hemisphere. The injector cannula remained in place for 1-minute post infusion to allow for liquid diffusion. The injector was then removed, and a sterilized dummy cannula was secured into the guide cannula. Inactivation was achieved by infusing Lidocaine hydrochloride (Sigma-Aldrich) diluted to 4% w/v in phosphate-buffered saline (PBS). The concentration and volume were selected to match those used by others to induce behavioral deficits in other learning paradigms (Lopez et al., 2012; Chang & Laing., 2017). Given the 0.5 ul infused here, a functional spread of slightly less than 1.0 mm would be expected (Martin, 1991; Welsh and Harvey, 1991). The short duration of the test phase (~ 1-3 min), run ~7 min after the infusion, fell within the window of lidocaine effect (Malpeli et al., 1999). The control condition, designed to control for the influences of the infusion process, was a volume matched infusion of PBS. For each animal, two lidocaine and two PBS infusions were performed for the 8 min retention interval trials examined here.

### 2.5. Histology

After completing all trails, rats were sacrificed via isoflurane overdose and perfused intracardially with saline followed by a 10% formalin solution. Brains were extracted and stored in formalin. 72 hours prior to slicing, brains were transferred into a 30% sucrose solution. Brains were sectioned at 40 um along the coronal plane and stained with 0.5% cresyl violet. Stained slices were imaged, and cannulas tip placement was confirmed by reference to a rat brain atlas as shown in Figure 1C (Paxinos & Watson 2007).

### 2.6. Data Analysis & Statistics

Memory performance at test was scored by 1) Percent correct: the percentage of the first four arms visited that were baited, and 2) Number of arm entries: the number of arms visited by the time the fourth correct arm was visited. An arm visit occurred when all four feet of the rat where in the arm. Statistical analysis for within group comparisons on both percent correct and total arm entry measures were performed with one-way repeated measures ANOVAs with two levels (PBS, Lidocaine), all p-values reported are Wilk’s Lambda test. Independent samples t-tests with two groups (cohort 1 vs cohort 2) were run to compare number of days to reach criterion first time, and number of days to return to criterion post-surgery prior to the start of infusions. Significance was defined at the α = 0.05 level. Reported values indicate mean ± standard deviation.

## 3. Results

To test whether the dorsal hippocampus is necessary for accurate test performance in a delayed win-shift spatial working memory task, we compared the percent accuracy and number of arm entries at test in animals that received lidocaine or PBS infusions between the study and test phases.

Performance did not significantly differ between the lidocaine and PBS conditions in cohort one whether it was measured as the percentage of the first four arm entries that were correct (81 ± 14% vs. 83 ± 17%, Wilk’s λ = 0.98, F(1,17) = 0.39 p = 0.54; Figure 2A) or as the total number of arm entries needed to collect all four rewards (5.2 ± 1.1 arms vs. 4.8 ± 0.9 arms, Wilk’s λ = .89, F(1,17) = 2.07, p = 0.17; Figure 2B). That is, rats in cohort one performed well whether or not their dorsal hippocampus was inactivated prior to test. The lack of difference between these conditions suggests that the accurate performance of the radial arm maze DSWS task does not always necessitate the dorsal hippocampus.

**Fig 2.**
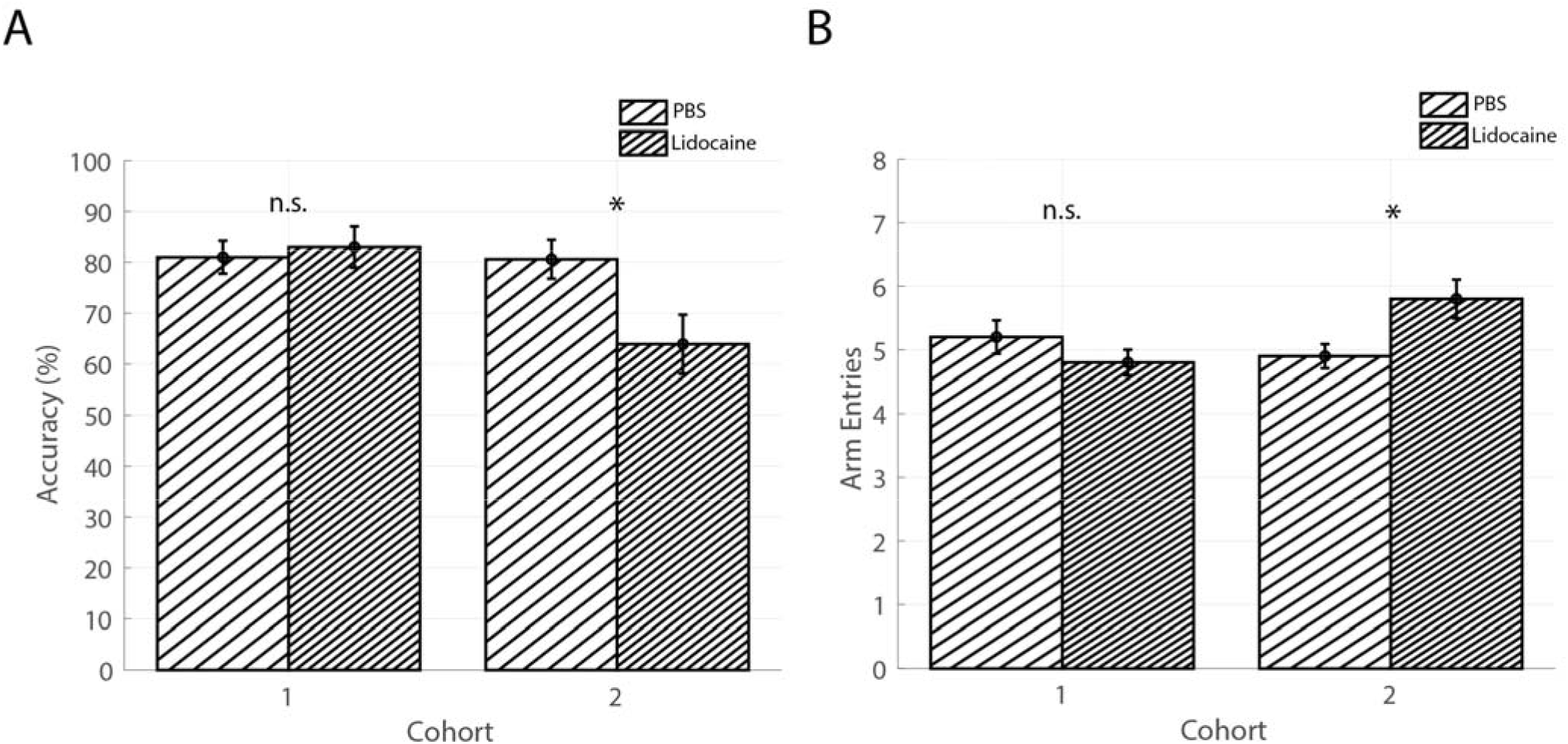
Dorsal hippocampal inactivation only impaired DSWS performance in cohort two. (A) Percentage of first four arm entries that were correct for both cohorts. No difference was observed in cohort one. In cohort two, the accuracy was significantly lower following lidocaine infusions. (B) Total arm entries to visit all four target arms for both cohorts. No difference was observed in cohort one. In cohort two, the number increased significantly following lidocaine infusions. Bar height reflects means, error bars indicate ±SEM. * - p < 0.05; n.s. – not significant.

Prior studies strongly suggest that the dorsal hippocampus should be required for this task. We hypothesized that the lack of hippocampal dependence observed in the first cohort was due to over training. Between reaching behavioral criterion for the first time and receiving the cannulation surgery, rats in the first cohort participated in two pilot experiments. This extending the amount of task training they received by an average of 28.3 ± 8.1 days. To test whether less training would uncover a sensitivity to the hippocampal inactivation, we ran a second cohort of rats. These received an average of only 7.6 ± 4.5 days of training between reaching criterion and undergoing surgery, differing significantly from the first cohort (F(1,16) = 4.49, p = < 0.0001). Consistent with the idea that the first cohort had a better grasp of the task prior to surgery, they returned to criterion significantly faster than the second cohort following surgery (6.4 ± 3.2 days vs. 17.3 ± 11.0 days, F(1.16) = 3.68, p = 0.011). Importantly, however, both cohorts reached the initial criterion (before surgery) in a similar number of days (17.44 ± 9.54 days vs. 23.44 ± 15.6 days, F(1,16) = 3.383, p = .340), demonstrating that there was not a systematic difference in capability between the cohorts. It is also important to note that in performing the two pilot experiments, the first cohort also received experience (12.55 trials on average) with retention intervals of between 60 and 150 minutes whereas the second cohort remained naïve to these long delays.

Consistent with the hypothesis that the additional experience the first cohort received reduced the necessity of the hippocampus for accurate performance at test, the second cohort, which did not receive this experience, was sensitive to hippocampal inactivation. Test performance in the second cohort was significantly lower following lidocaine infusions relative to PBS infusions whether measured by percent correct (81 ± 16% vs. 64 ± 25%, Wilk’s λ = 0.667, F(1,17) = 8.50, p = 0.010; Figure 2A) or by arm entries (4.9 ± 0.8 arms vs. 5.8 ± 1.3 arms, Wilk’s λ = .704, F(1,17) = 7.15, p = 0.016; Figure 2B). The sensitivity of the second cohort to hippocampal inactivation demonstrates that the dorsal hippocampus is involved performance of the delayed spatial win-shift task, at least initially. When considered together with the data from the first cohort, the present findings suggest that, with sufficient training, performance of this task become independent of the dorsal hippocampus.

## 4. Discussion

Understanding the brain structures that play a necessary role in spatial working memory remains an outstanding challenge. The traditional view holds that, when retention intervals span minutes in a delayed spatial win shift (DSWS) task and the arms shown at study are randomized over trials, the hippocampus is always required. In this brief report, we describe data showing that radial arm maze DSWS task performance was not impaired when lidocaine was infused into the dorsal hippocampus prior to the test phase in a cohort of rats. The same infusions did impair performance, however, in a second cohort of rats that had received significantly less task training. These results indicate that the dorsal hippocampus is important for DSWS task performance at test early in training but, with additional experience, becomes less important.

Our finding that lidocaine infusions did not impair performance when done prior to the 8 min tests was surprising given previous work showing the importance of the hippocampus for working memory at similar delays (Lee & Kesner., 2003a; Lee & Kesner., 2003b; Yoon et al., 2008; Churchwell & Kesner., 2011). Lee & Kesner (2003a), for example, showed that temporary inactivation/lesion of dorsal hippocampus/mPFC during a radial arm DSWS task impaired performance with a 5-minute retention interval. Importantly, the deficit could not be compensated for by an intact mPFC indicating the specific importance of the dorsal hippocampus at such delays.

The discrepancy between prior reports and our findings may be attributable to methodological differences. The most significant difference between prior studies and ours is that in prior work the inactivation/lesions had effect on both the study and test phases. Here, the inactivation was performed between the study and test phases leaving the dorsal hippocampus fully functional during the study phase. As such, unlike prior work, the manipulation used here asked whether the dorsal hippocampus is necessary specifically at test. Unfortunately, we were not able to test this hypothesis directly by performing the inactivation prior to the study phase in the first cohort because this striking pattern of results was identified incidentally in analysis of data collected as a control for a larger study after the rats had already been sacrificed. Future research will be needed to test this hypothesis directly.

Another key difference may be the amount of training the animals received. The cohort of rats that was insensitive to the inactivation had received considerably more training on the task prior to the cannulation surgery. With this training, they also received experience with delays of 60 minutes or longer. Given the cohort that did not receive the extra training or experience with long delays remained sensitive to the inactivation, it is likely that this experience contributed to the reduction in hippocampal dependence. Whether it was the extra trials of experience or specifically the experience with the long delays that led to this shift remains unclear. The experimental design used here was poorly suited to sorting between these possibilities because it was initially designed for a separate study.

In summary, we describe here the incidental finding that pharmacological inactivation of the dorsal hippocampus prior to the test phase of a DSWS task did not impair spatial working memory performance in a cohort of rats. While in a second cohort, the task was sensitive to dorsal hippocampus inactivation. This finding is striking in that it contrasts with what prior work has shown on hippocampal contributions to spatial working memory would have predicted. These results do not directly contradict prior findings however, as there are important procedural differences between what was done here and what has been done previously. Perhaps most importantly, this is the first study to test the effect of dorsal hippocampal inactivation selectively during the DSWS test phase. The present results were found during analysis of non-essential trials of a larger study and additional work is needed, using purpose designed experiments, to better understand the boundary conditions of the current findings.

## Acknowledgements

This work was made possible by generous funding from the Harlan Scholars Program and the IU-MSI STEM Initiative. We graciously thank the Indiana University Laboratory Animal Resources facilities for their attention and care of our animals.

## References

Buzsaki G., & Moser E. I. (2013). Memory, navigation and theta rhythm in the hippocampal-entorhinal system. Nature Neuroscience, 16, 2, 130–8.

Chang S. D., & Liang K. C. (2017). The hippocampus integrates context and shock into a configural memory in contextual fear conditioning. Hippocampus, 27, 2, 145–155.

Churchwell J. C., & Kesner R. P. (2011). Hippocampal-prefrontal dynamics in spatial working memory: interactions and independent parallel processing. Behavioural Brain Research, 225, 2, 389–95.

Crystal J. D., Ketzenberger J. A., & Alford W. T. (2013). Practicing memory retrieval improves long-term retention in rats. Current Biology: Cb, 23, 17, 708–9.

Floresco S. B., Seamans J. K., & Phillips A. G. (1997). Selective roles for hippocampal, prefrontal cortical, and ventral striatal circuits in radial-arm maze tasks with or without a delay. The Journal of Neuroscience: the Official Journal of the Society for Neuroscience, 17, 5, 1880–90.

Hartley T., Lever C., Burgess N., & O’Keefe J. (2014). Space in the brain: how the hippocampal formation supports spatial cognition. Philosophical Transactions of the Royal Society of London. Series B, Biological Sciences, 369, 1635.).

Lee I., & Kesner R. P. (2003a). Time-dependent relationship between the dorsal hippocampus and the prefrontal cortex in spatial memory. The Journal of Neuroscience: the Official Journal of the Society for Neuroscience, 23, 4–1517.

Lee I., & Kesner R. P. (2003b). Differential roles of dorsal hippocampal subregions in spatial working memory with short versus intermediate delay. Behavioral Neuroscience, 117(5), 1044–1053.

Lopez J., Herbeaux K., Cosquer B., Engeln M., Muller C., Lazarus C., Kelche C., … de V. A. P. (2012). Context-dependent modulation of hippocampal and cortical recruitment during remote spatial memory retrieval. Hippocampus, 22, 4, 827–841.

Malpeli J. G. (1999). Reversible inactivation of subcortical sites by drug injection. Journal of Neuroscience Methods, 86, 2, 119–128.

Martin J. H. (1991). Autoradiographic estimation of the extent of reversible inactivation produced by microinjection of lidocaine and muscimol in the rat. Neuroscience Letters, 127, 2, 160–164. Bottom of Form.

McDaniel W. F., Compton D. M., & Smith S. R. (1994). Spatial learning following posterior parietal or hippocampal lesions. Neuroreport, 5, 14, 1713–1717.

O’Keefe J., & Nadel L. (1978). The hippocampus as a cognitive map. Oxford: Clarendon Press.

Olton DS, Samuelson RJ (1976) Remembrance of places passed: Spatial memory in rats. Journal of Experimental Psychology: Animal Behavior Processes 2: 97–116.

Packard M. G., Regenold W., Quirion R., & White N. M. (1990). Post-training injection of the acetylcholine M 2 receptor antagonist AF-DX 116 improves memory. Brain Research, 524, 1, 72–76.

Paxinos G., & Watson C. (2007). The rat brain in stereotaxic coordinates. Amsterdam: Elsevier.

Phillips A. G., Ahn S., & Floresco S. B. (2004). Magnitude of dopamine release in medial prefrontal cortex predicts accuracy of memory on a delayed response task. The Journal of Neuroscience: The Official Journal of the Society for Neuroscience, 24, 2, 547–53.

Potvin O., Allen K., Thibaudeau G., Dore F. Y., & Goulet S. (2006). Performance on spatial working memory tasks after dorsal or ventral hippocampal lesions and adjacent damage to the subiculum. Behavioral Neuroscience, 120, 2, 413–22.

Seamans JK, Phillips AG (1994) Selective memory impairments produced by transient lidocaine-induced lesions of the nucleus accumbens in rats. Behav Neurosci 108:456–468. Bottom of Form.

Seamans JK, Floresco SB, Phillips AG (1995) Functional differences between the prelimbic and anterior cingulate regions of the rat prefrontal cortex. Behavioral Neuroscience 109: 1063–1073.

Taylor C. L., Latimer M. P., & Winn P. (2004). Impaired delayed spatial win-shift behaviour on the eight arm radial maze following excitotoxic lesions of the medial prefrontal cortex in the rat. Behavioural Brain Research, 147, 1, 107.

Welsh JP, Harvey JA (1991) Pavlovian conditioning in the rabbit during inactivation of the interpositus nucleus. J Physiol. (Lond) 444:459–480.

Yoon T., Okada J., Jung M. W., & Kim J. J. (2008). Prefrontal cortex and hippocampus subserve different components of working memory in rats. Learning & Memory, 15, 3, 97–105.

